# Noise-induced scaling in skull suture interdigitation

**DOI:** 10.1101/2020.06.24.168674

**Authors:** Yutoh Naroda, Yoshie Endo, Kenji Yoshimura, Hiroshi Ishii, Shin-Ichiro Ei, Takashi Miura

## Abstract

Sutures, the thin, soft tissue between skull bones, serve as the major craniofacial growth centers during postnatal development. In a newborn skull, the sutures are straight; however, as the skull develops, the sutures wind dynamically to form an interdigitation pattern. Moreover, the final winding pattern had been shown to have fractal characteristics. Although various molecules involved in suture development have been identified, the mechanism underlying the pattern formation remains unknown. In a previous study, we reproduced the formation of the interdigitation pattern in a mathematical model combining an interface equation and a convolution kernel. However, the generated pattern had a specific characteristic length, and the model was unable to produce a fractal structure with the model.

In the present study, we focused on the anterior part of the sagittal suture and formulated a new mathematical model with time–space-dependent noise that was able to generate the fractal structure. We reduced our previous model to represent the linear dynamics of the centerline of the suture tissue and included a time–space-dependent noise term. We showed theoretically that the final pattern from the model follows a scaling law due to the scaling of the dispersion relation in the full model, which we confirmed numerically. Furthermore, we observed experimentally that stochastic fluctuation of the osteogenic signal exists in the developing skull, and found that actual suture patterns followed a scaling law similar to that of the theoretical prediction.

**Author summary:** Skull sutures (thin, undifferentiated tissue between bones) act as the growth centers for the skull. Sutures are straight at birth but later develop an interdigitated pattern that ultimately becomes a fractal structure. While our previous mathematical model of sutures generated a periodic pattern, the mechanism underlying the fractal structure formation remained to be elucidated. Here, we focused only on the anterior part of the sagittal suture and formulated a reduced model representing the initial linear phase of pattern formation with the addition of a time–space-dependent noise term. We showed analytically that the model generates patterns with a scaling law. This result was confirmed numerically and experimentally.

## Introduction

Sutures are the thin, soft tissues between skull bones. They perform multiple functions, and they have been extensively studied as a model system of skeletal development [1]. During development, the suture tissue acts as the growth center of the skull; the premature disappearance of the suture tissue causes the skull deformation known as craniosynostosis [2]. At birth, the suture tissue is thick and straight; however, as the skull develops, the sutures gradually becomes thinner and begins to form winding and interdigitated patterns [3]. After adolescence, the skull bones gradually fuse and the suture tissues disappear. Consequently, suture tissue can be used to estimate the age of a person in forensic science techniques [4]. The suture tissue is known to mechanically connect the skull bones, and the interdigitation is assumed to reinforce the mechanical strength of the connection [5].

The interdigitation of sutures results in a fractal structure, which was initially reported in the mid-1980s [6,7]. Until recently, researchers generally measured the fractal dimension by the box-counting method [8–13]. These measurements have mainly been used for classification or diagnostic purposes; however, studies are yet to reveal the mechanism of fractal pattern formation.

Although various models have been proposed for the formation of fractal structures, e.g., the Eden collision model [14], the Koch curve [15] and diffusion-limited aggregation [16], their application to suture pattern formation has been unsuccessful. For example, (i) Oota et al. [17, 18] applied the Eden model to the formation of the curvature of skull sutures, (ii) Zollikofer and Weissmann applied the diffusion-limited aggregation method to the formation of skull suture interdigitation [16], and (iii) we suggested that the formation of skull suture curvature based on the time-dependent diffusion term was equivalent to the Koch curvature generation [3]. However, these models produced results inconsistent with the experimental observations. For example, in our previous study [3], we introduced time-dependent diffusion terms to reproduce the fractal structure [3]; however, in a subsequent study that used computed tomography, we were unable to detect the addition of small structures to the large structures during development, which is expected in a model with time-dependent diffusion terms [19].

In the present study, therefore, we propose a new mechanism for suture fractal pattern formation that incorporates a noise term. We focus on the anterior part of the sagittal suture, which exhibits more random characteristics than other sutures (Fig. 1). In order to analyze this mathematically, we reduced our previous model [19] to the dynamics of the centerline *h*(*x,t*). Subsequently, we introduce time-space-dependent noise to the model and show that such a system with noise can exhibit scaling. Finally, we demonstrated experimentally that the noise of the osteogenic signal existed in the developing skull, and the scaling of the anterior part of the sagittal suture was consistent with the model predictions. The overall structure of this study is schematically presented in Fig. 2.

**Fig 1.**
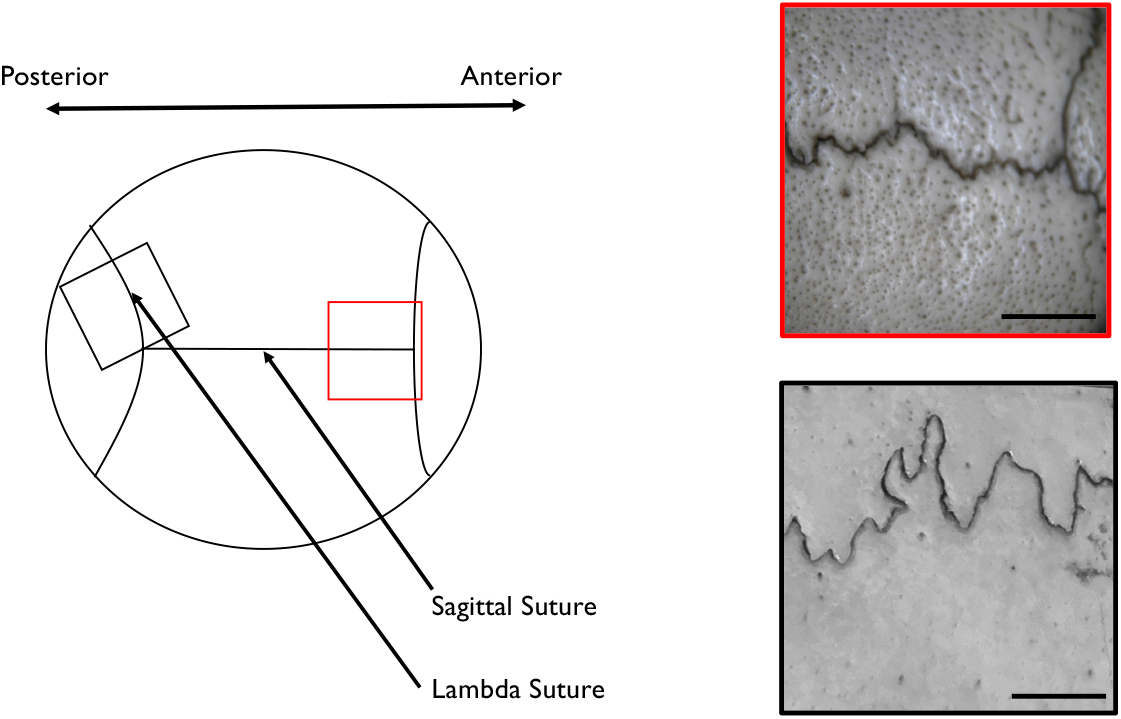
The anterior part of the sagittal suture in an adult skull shows stochastic small amplitude interdigitation. The suture in the anterior skull near the bregma (red box) has a rough surface with less prominent curvature and a more stochastic appearance than that of the lambda suture (black box). Scale bars (bottom right of boxes) = 0.5 cm.

**Fig 2.**
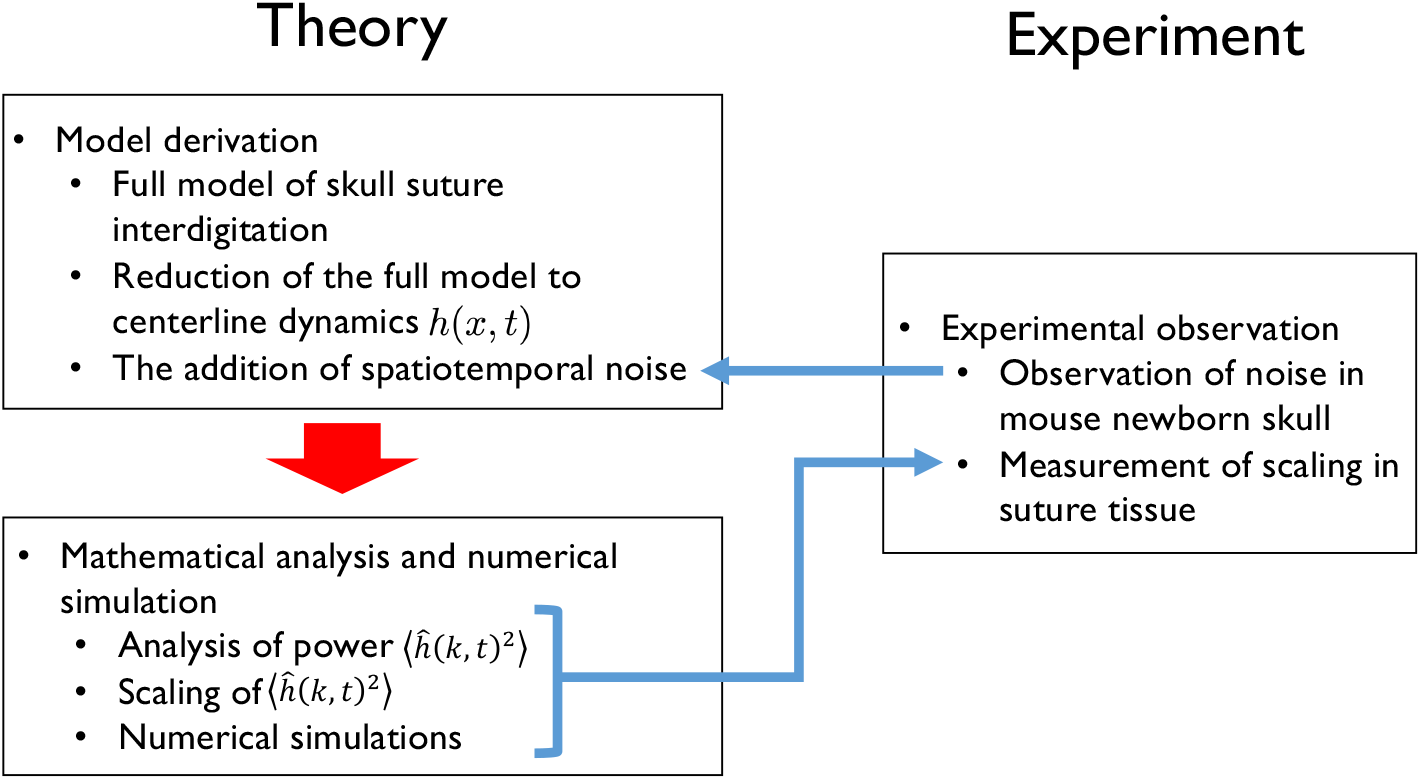
Schematic diagram of the present study. First, we reduced our previous model [19] into graph form *h*(*x, t*). Next, we added a noise term to the model based on our experimental observation of noise (Fig. 6). Then we mathematically derived the scaling law from this equation and confirmed it using numerical simulations. Finally, we showed experimentally that both noise and scaling do exist in skull sutures (Fig. 7).

## Materials and methods

### Observation of noise in newborn mouse skull

To observe spatiotemporal noise in a developing mouse skull, we used a transgenic mouse that expressed a FRET sensor for the phosphorylated ERK signal [20] as well as an organ culture system. We sacrificed newborn mice by decapitation and dissected their skulls using forceps and a scalpel. The isolated skulls were then placed onto Millicell culture inserts (Millipore Inc.) with a Dulbecco’s modified Eagle medium/F-12 culture medium containing 10% fetal bovine serum and antibiotics. We observed the cyan and yellow fluorescent protein ratio (CFP/YFP ratio) using a Nikon A1R confocal microscope, and we quantified the spatiotemporal fluctuation using Fiji software [21]. This experiment was undertaken with the permission of the Kyushu University animal experiment committee (A29-036-1).

### Measurements of sagittal sutures

Anterior sagittal skull sutures in human bone specimens from Kyushu University School of Medicine and School of Dentistry were digitized using a Stemi 2000-CS stereomicroscope (0.65x magnification; Carl Zeiss) with a digital camera (COOLPIX P6000; Nikon) and an adapter (NY-P6000 Super; Microscope Network Co., Ltd.). Detailed information about these skulls, including their origins, was not available.

We took photographs of the sagittal suture at a distance of 2 cm from the glabella. We then selected 14 sutures with small amplitude interdigitation and without overhang, and we printed images of these sutures onto A4-sized sheets of paper with reduced contrast. For each image, we traced the sagittal suture with a red marker pen and used a document scanner (ScanSnap iX500; Fujitsu) to scan the image into a computer. Using Fiji software [21], we first separated the red channel of the scan to obtain the suture line, which was then skeletonized in order to detect its x and y coordinates. Finally, the coordinate data were mathematically analyzed using *Mathematica* (Wolfram Research).

## Results

### Model derivation

#### Full model of skull suture interdigitation

We previously proposed the following simple model of skull suture interdigitation [19]:

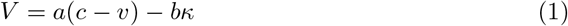

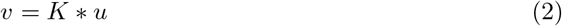

where *V* is the interface speed perpendicular to the bone–mesenchyme interface, *a* is the efficiency of the substrate factor (the osteogenesis-promoting diffusible signaling molecules expressed at the mesenchyme) over bone differentiation, c is the threshold value for bone generation/resorption. *b* is the surface tension, and *κ* is the local curvature. In addition, *v*(*x,y,t*) represents the effect of the substrate factor, determined by the convolution of the kernel *K*(*x,y*) and bone shape *u*(*x,y,t*) (where *u*(*x,y,t*) = 1 represent bone and *u*(*x,y,t*) = 0 represent mesenchyme). This system has a band-like solution with interface instability, and its linear dynamics are well understood (see [19] and subsection A. in S1 text). With a certain parameter set, the real part of the eigenvalue λ(*k*) takes a maximum positive value at *k*_max_, indicating the emergence of a structure with a specific wavenumber (Fig. 8 in S1 text). However, with this model, the fractal structure should not appear.

In the present study, we focused on sections of sutures with less pronounced curvature (red panel in Fig. 1). Therefore we used the parameter sets that satisfy λ(*k*) < 0 for all *k* (lower left part of Fig. 8 in S1 text).

#### Reduction of the full model to centerline dynamics *h*(*x, t*)

Next, we further simplified the model to represent only the linear dynamics of the full system as follows (Fig. 3):

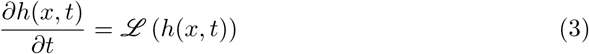

where *h*(*x,t*) is the y-coordinate of the suture at x-coordinate *x* and time *t*. Only the small amplitude patterns without overhangs were considered. Here, *ℒ* represents a linear operator that reproduces the linear dynamics of the full model given by equations (1) and (2). The explicit form of *ℒ* is described in subsection B. in S1 text. Thus, the Fourier transformation of *ℒ* should be *λ*(*k*) given by equation (14) (subsection A. in S1 text), and the frequency domain for the system (equation (3)) is as follows:

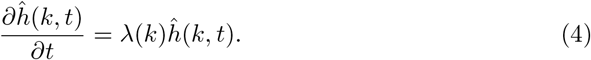

**Fig 3.**
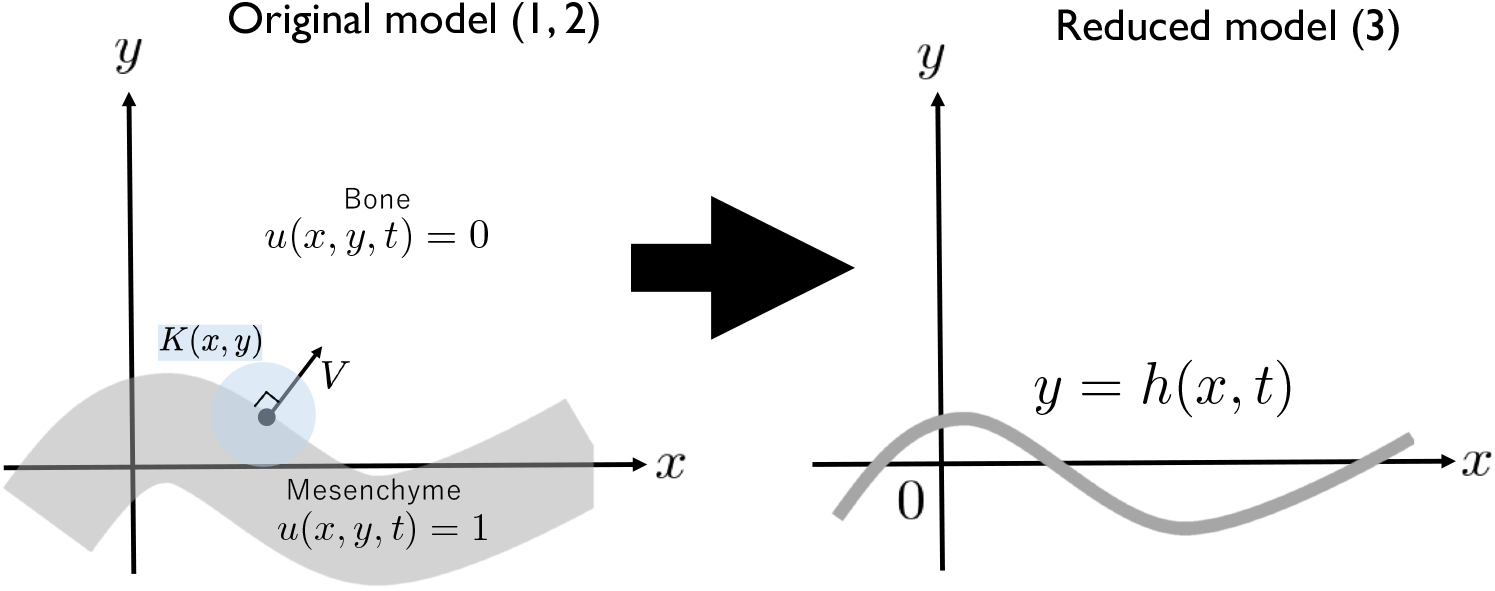
Model reduction. The original model (equations (1, 2)) considered a band-like solution with width 2*y*_0_. The present model (equation (3)) focuses on the dynamics of the centerline *h*(*x, t*) of the band-like solution, considering only the onset of pattern formation without overhangs.

#### The addition of spatiotemporal noise

In this section, we incorporated the noise term *H* in the full system (1,2) as follows:

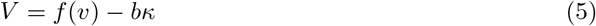

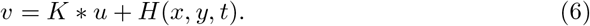

Here, *H* was defined as the spatiotemporal white noise caused by the known stochastic fluctuation of gene expression ([22], Fig. 6).

**Fig 4.**
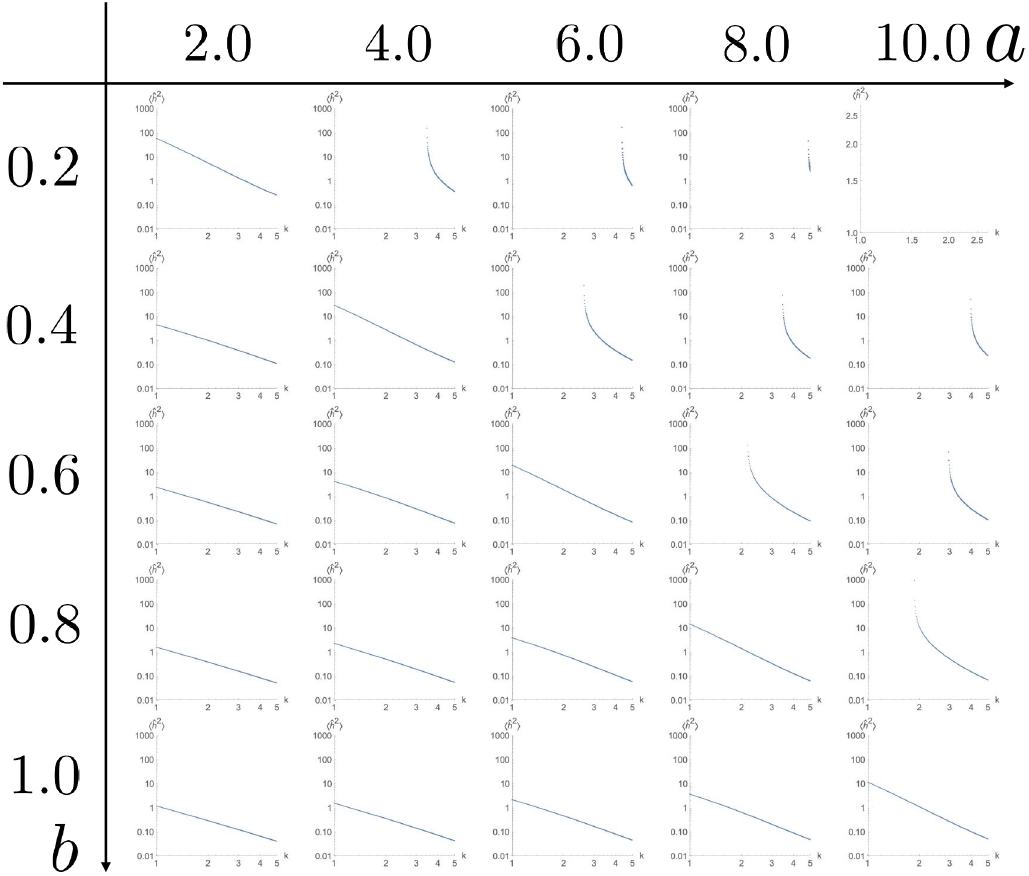
Log–log plot of the (*k*, –1/λ(*k*)) of full model, which should reflect the power spectrum 〈*ĥ*(*k, t*)^2^〉. The distribution shows linearity in the parameter range without spontaneous pattern formation, indicating λ = –*γk^β^*.

**Fig 5.**
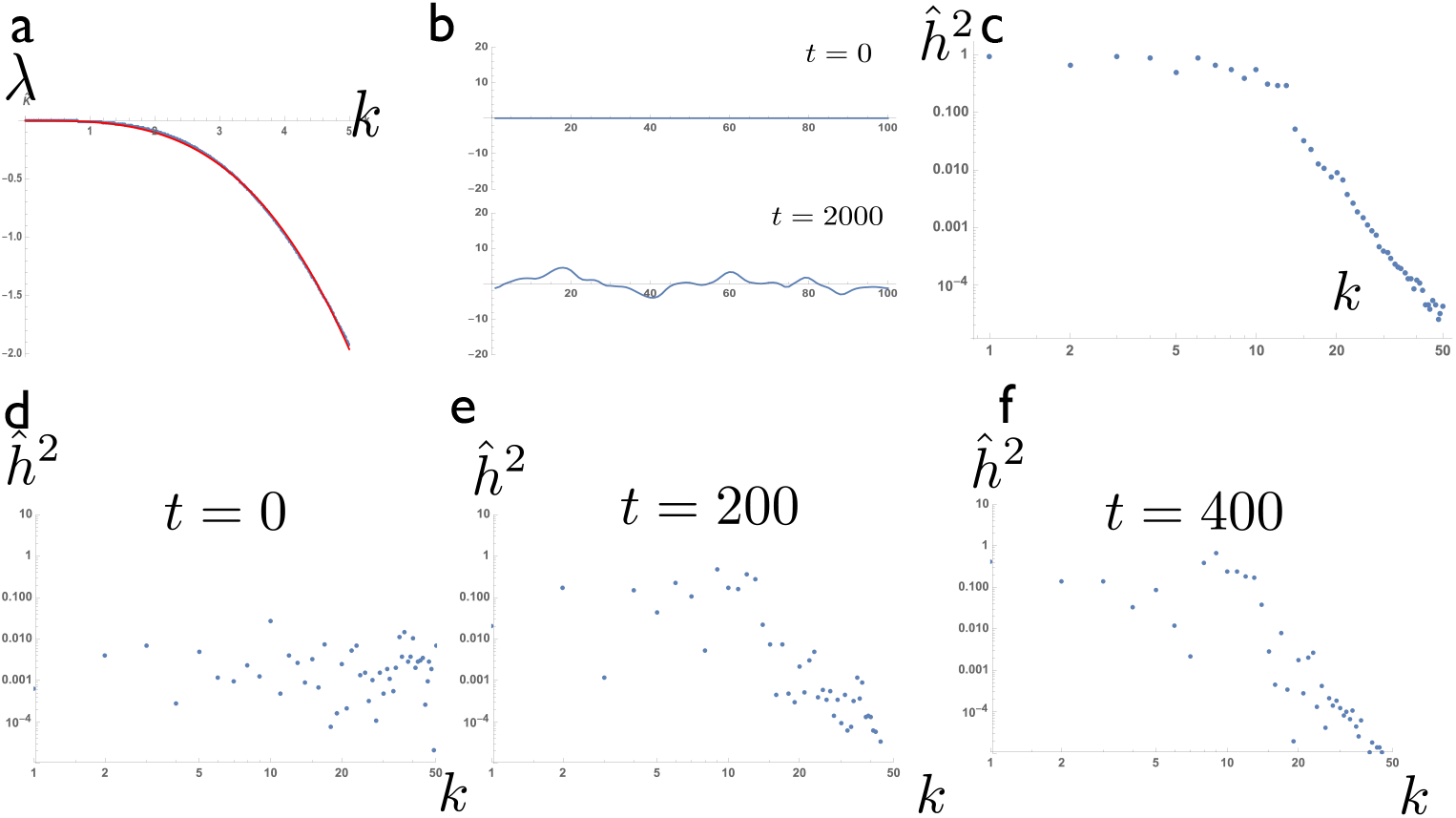
Numerical simulation of the reduced model showed the scaling predicted by the mathematical analysis. (a) Dispersion relation of the full model (blue) and its approximation by λ(*k*) = –*γk^β^* (red). The parameter set was *a* = 1, *b* = 0.1, *c* = 0.48, and *r* =1. (b) Result of the numerical simulation. The upper panel shows the initial shape (*t* = 0); the lower panel shows the curved shape obtained after sufficient time had passed (*t* = 2000). (c) Log–log plots of the average of *ĥ*^2^. A region of linear scaling was observed in the high wavenumber region. (d–f) Time course of the log-log plots of the power spectrum. (d) *t* = 0, (e) *t* = 200, and (f) *t* = 400.

**Fig 6.**
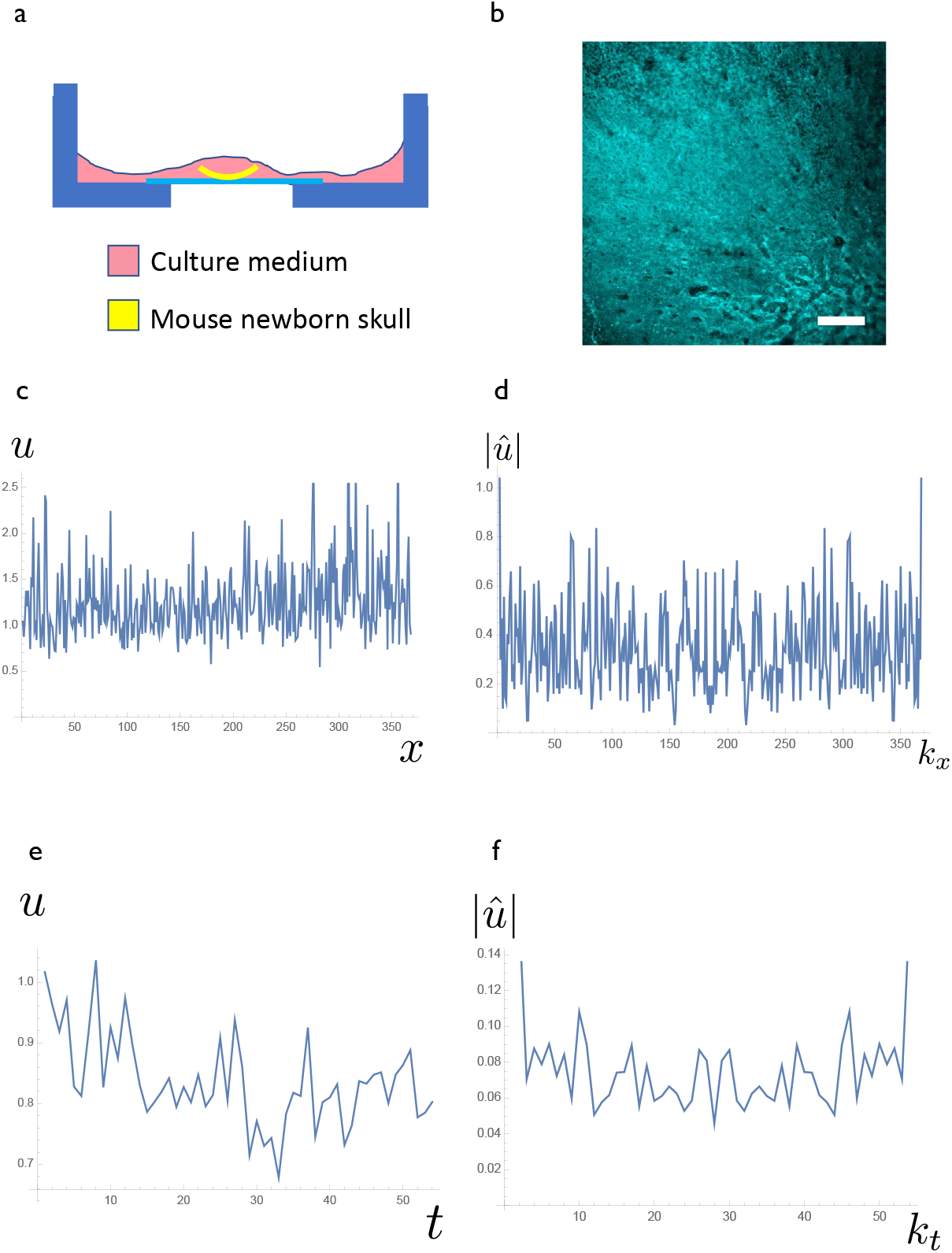
Direct observation of the intrtinsic noise of the osteogenic signal in a newborn mouse skull. (a) Experimental system setup. A newborn skull of an ERK-FRET transgenic mice was dissected and then set on the glass bottom dish, which was suitable for organ culture. (b) Image of the cyan and yellow fluorescent protein ratio (CFP/YFP ratio) representing the ERK signal input. The signal in the skull was spatially heterogeneous. Scale bar = 200 *μm*. (c) Spatial distribution of the FRET signal *u*. (d) Fourier transformation of *u*(*x, t*). There was no specific characteristic in the distribution, indicating the noise was white noise. The distribution contained no specific characteristics, indicating that the noise was white noise. (e) Time course of FRET signal intensity at a specific point. The signal showed fluctuation. (f) Fourier transformation of (e). No specific trend was observed, indicating that the spatial noise could be regarded as white noise.

In the full model, the band-like solution moves according to the *gradient* of the noise 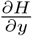, not according to the magnitude of the noise itself (subsection D. in S1 text). However, the reduced model considers only changes with small amplitudes without overhangs, neglecting movement in the *x*-direction, so it can be assumed that the noise term at the suture point (*x, h*(*x, t*)) is white noise (Fig. 12 in S1 text). The final reduced model with spatiotemporal noise is as follows:

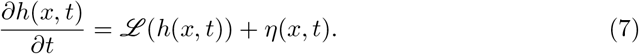

### Mathematical analysis and numerical simulation

#### Analysis of power 〈*ĥ*(*k, t*)^2^〉

In a fractal structure, the measurement scale and measured quantity have a linear relationship in log-log plots. For the frequency domain, the wavenumber *k* and power 〈*ĥ*(*k, t*)^2^〉 should show linearity on log–log plots. This implies that

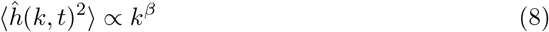

holds in a fractal structure; an intuitive explanation of this is presented in subsection C. in S1 text. The goal of this analysis was to show the relationship in equation (8).

We assume that *η*(*x,t*) is white noise with a mean value of 0 and variance *D*.

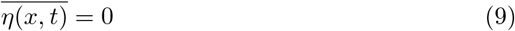

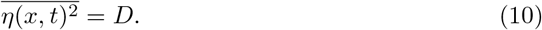

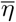 represents the spatial average of *η*.

The Fourier transformation of (3) is

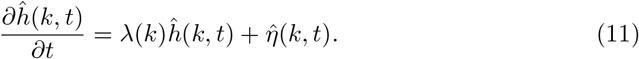

The steady-state sample mean of *ĥ*(*k,t*)^2^ can be obtained by taking the limit as *t* → ∞:

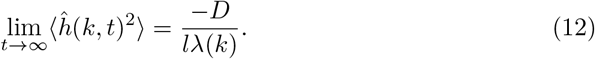

Detailed derivation is described in the subsection E. in S1 text. The sample mean of the *ĥ* variance 〈*ĥ*^2^〉 in the steady state (12) is inversely proportional to λ(*k*). Therefore, when λ(*k*) can be approximated by λ(*k*) ∝ *k^β^* at a certain spatial scale, the resulting pattern can be fractal.

#### Scaling of 〈*ĥ*(*k, t*)^2^〉

If we can approximate λ ∝ *k^β^*, then the resulting pattern has some scaling. In other words, the log-log plot of (*k*, –1/λ(*k*)) should show linearity, and thereby indicate that λ = –*γk^β^*. *γ* is a positive proportionality constant.

We systematically changed the model parameter set (*a, b*), and construct log–log plots of (*k*, –1/λ(*k*)). In very large and small spatial scales, scaling was determined by the surface tension term (subsection F. in S1 text). However, in some parameter regions, the plots showed linearity within a certain range of spatial scale (Fig. 4, lower left half), indicating that a fractal structure should arise in this parameter region. We found that the gradient varied depending on the parameter set (*a, b*).

#### Numerical simulations

Numerical simulations of our model (equation (3))showed that the reduced model could generated patterns with scaling (Fig. 5). We initially chose a parameter set that did not exhibit interface instability (Fig. 5a). Linearity in log–log plots indicated that the dispersion relation in this range could be approximated by –*γk^β^*. Importantly, the numerical simulation produced a suture pattern that resembled the small amplitude interdigitation observed in vivo (Fig. 1, Fig. 5b). Because of the continuous input of noise *η*, the distribution did not reach a steady-state (Fig. 5c). However, the scaling expected from the mathematical analysis was achieved after a sufficiently long simulation time (Fig. 5d-f).

### Experimental observation

#### Observation of noise in a newborn mouse skull

We introduced noise term *η* in the governing equation (7). To investigate the characteristic of noise term, we observed ERK signal input fluctuation using ERK-FRET mice [20]. One of the major signals for osteogenesis is the FGF pathway [3], and the phosphorylation of ERK should represent the signal input of this FGF pathway [20]. We set up an organ culture system of the newborn mouse skull expressing ERK-FRET sensor (Fig. 6a) and observed the spatiotemporal signal fluctuation (Fig. 6b). We observed the spatial and temporal fluctuation of the ERK input signal (Fig. 6.a-d). The power spectrum of spatial and temporal noise seemed flat (Fig. 6.c-f), indicating that the observed noise was white noise as assumed in the model (7).

#### Measurement of scaling in human suture tissue

We obtained *β* from trace data from human sutures. We obtained coordinate data for the anterior part of a sagittal suture (Fig. 7a) and derived the power spectrum of the suture line; a log-log plots of the power spectrum showed linearity (Fig. 7b). The slope of the distribution was then obtained using the least-square method. This corresponded to –*β*. The mean and standard deviation of measured 2β were 2.8 and 0.4, respectively; our model (equation 7) was able to reproduce this scaling (Fig. 5).

**Fig 7.**
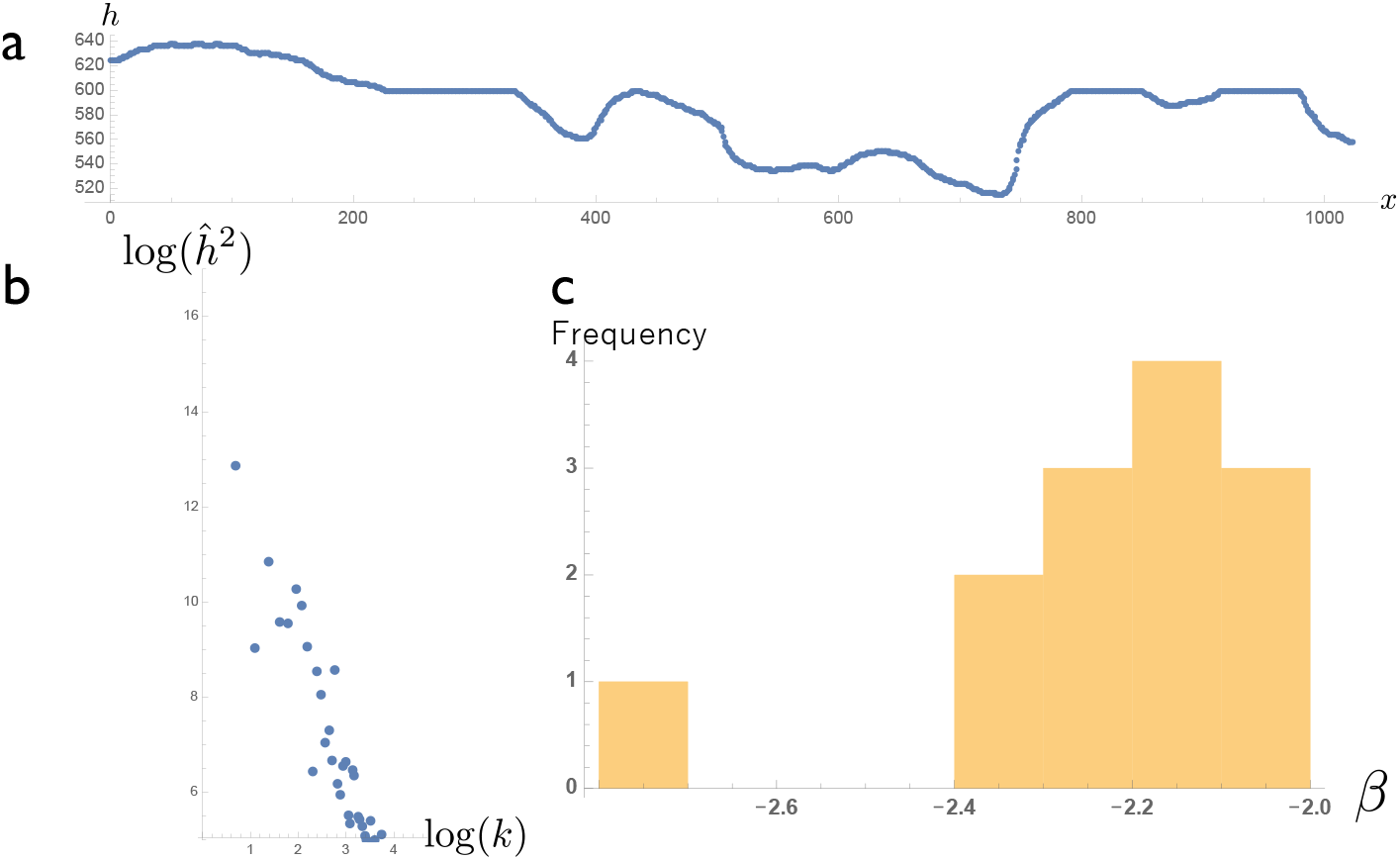
Scaling in actual skull sutures. (a) A traced suture line converted into coordinates and plotted on the *x–h* plane. (b) Relationship between *k* and *ĥ* on a log–log plot. (c) Histogram of *β*.

## Discussion

In this study, we formulated a simple model based on our previous work [19], and analytically obtained the scaling of the suture tissue, which was confirmed by experimental observations. Although many previous studies have investigated the fractality of skull sutures [8–12], ours is the first study in which the scaling observed in vivo has been directly correlated with a mathematical model of pattern formation. While previously published models have been able to generate a fractal structure, they were not fully consistent with experimental observations. For example, the Eden collision model [17,18]did not reproduce the initial phase of pattern formation, whereas the bidirectional growth model [16] generated structures that were too complex.

Moreover, neither of these previous models were analytically manageable.

In the present study, we obtained a scaling parameter *β*, which may reflect the pattern formation mechanism. However, the relationship between this scaling parameter and the previously measured fractal dimensions is unclear. We measured the fractal dimension of the suture by the divider method, but it produced a result close to 1, which means that *β* is not directly connected to the fractal dimension. However, due to its high sensitivity, *β* may be useful for diagnositic purposes. For example, in craniosynostosis, osteogenesis is promoted; in the full model, this means smaller c value than normal. In our analysis, *β* should become smaller by a small value of *c*. Therefore, smaller *β* in actual suture indicates a higher risk of craniosynostosis.

In future research, the goal will be to understand scaling in larger amplitude cases. In the present study, we dealt with scaling only for sutures with a small amplitude, but there are some types of suture curvature in which the amplitude of a specific frequency is emphasized. In such cases, the system exhibits interface instability, and a positive *λ_max_* exists, and the curvatures are much more pronounced. Our model cannot measure the scaling of sutures with such pronounced curvature and overhanging (such as lambda sutures) and can be applied only to small amplitude sutures.

Whether the fractal nature of suture tissue has a biological function remains to be elucidated. It has been postulated that the interdigitation strengthens the junction between the skull bones [5]. The functional difference between simple geometry (sine curves) and fractal structure has, however, been examined from an engineering perspective [23–25]. Here, we assessed the function of the fractal structure from a biological perspective by assuming that junction strength was correlated with the length of the junction (see subsection G. and H. in S1 text). However, we did not observe any clear increase in junction strength with the fractal structure. Although large-scale measurement of human suture strength has been done [26], the effect of interdigitation on suture strength remains unclear because the fusion of suture tissue may also influence its mechanical properties. A literature survey did not find any clear correlation between the degree of suture interdigitation and the prevalence of fractures [27]. Further study is necessary to fully understand the biological importance of the fractal nature of skull sutures.

## Supporting information

**S1 Text. Details of the models and mathematical analyses.**

## Ethics

This work was approved by Kyushu University Institutional Review Board for Clinical Research (2019-350). Human skull samples used in this study (n=50) were those used for the education of osteology at the Faculty of Medicine, Kyushu University. Since the samples were collected several decades ago and already anonymized, we could not obtain informed consent in this case. Therefore, we notified the general publifc of study via our laboratory homepage (http://www.lab.med.kyushu-u.ac.jp/anat1/) according to the rules of the local ethics committee.

## Acknowledgements

We want to thank Professor Toshio Kukita in the Department of Dentistry, Kyushu University, for providing the skull specimens, Nobuhide Shibusawa in Kyushu University for helping with skull observations, and Toshiki Oguma in Kyushu University, and Dr. Katsuhiko Sato and Yasuaki Kobayashi in Hokkaido University, for helpful discussions and comments. This work was financially supported by JSPS KAKENHI Grant Number 15KT0018.

